# Balancing limited resources in actin networks competition

**DOI:** 10.1101/2024.09.09.612098

**Authors:** Christophe Guérin, Anne-Betty N’Diaye, Laurène Gressin, Alex Mogilner, Manuel Théry, Laurent Blanchoin, Alexandra Colin

## Abstract

In cells, multiple actin networks coexist in a dynamic manner. These networks compete for a common pool of actin monomers and actin-binding proteins. Interestingly, this competition does not result in the mere survival of the more consuming networks. Moreover, the co-existence of networks with various strengths is key to cell adaption to external changes. However, a comprehensive view of how these networks coexist in this competitive environment, where resources are limited, is still lacking. To address this question, we used a reconstituted system, in closed microwells, consisting of beads propelled by actin polymerization or micropatterns functionalized with lipids capable of initiating polymerization close to a membrane. This system enabled us to build dynamic actin architectures, competing for a limited pool of proteins, over a period of hours. We demonstrated the importance of protein turnover for the coexistence of actin networks, showing it ensures resource distribution between weak and strong networks. However, when competition becomes too intense, turnover alone is insufficient, leading to a selection process that favors the strongest networks. Consequently, we emphasize the importance of competition strength, which is defined by the turnover rate, the amount of available protein, and the number of competing structures. More generally, this work illustrates how turnover allows biological populations with various competition strengths to coexist despite resource constraints.

Graphical Abstract

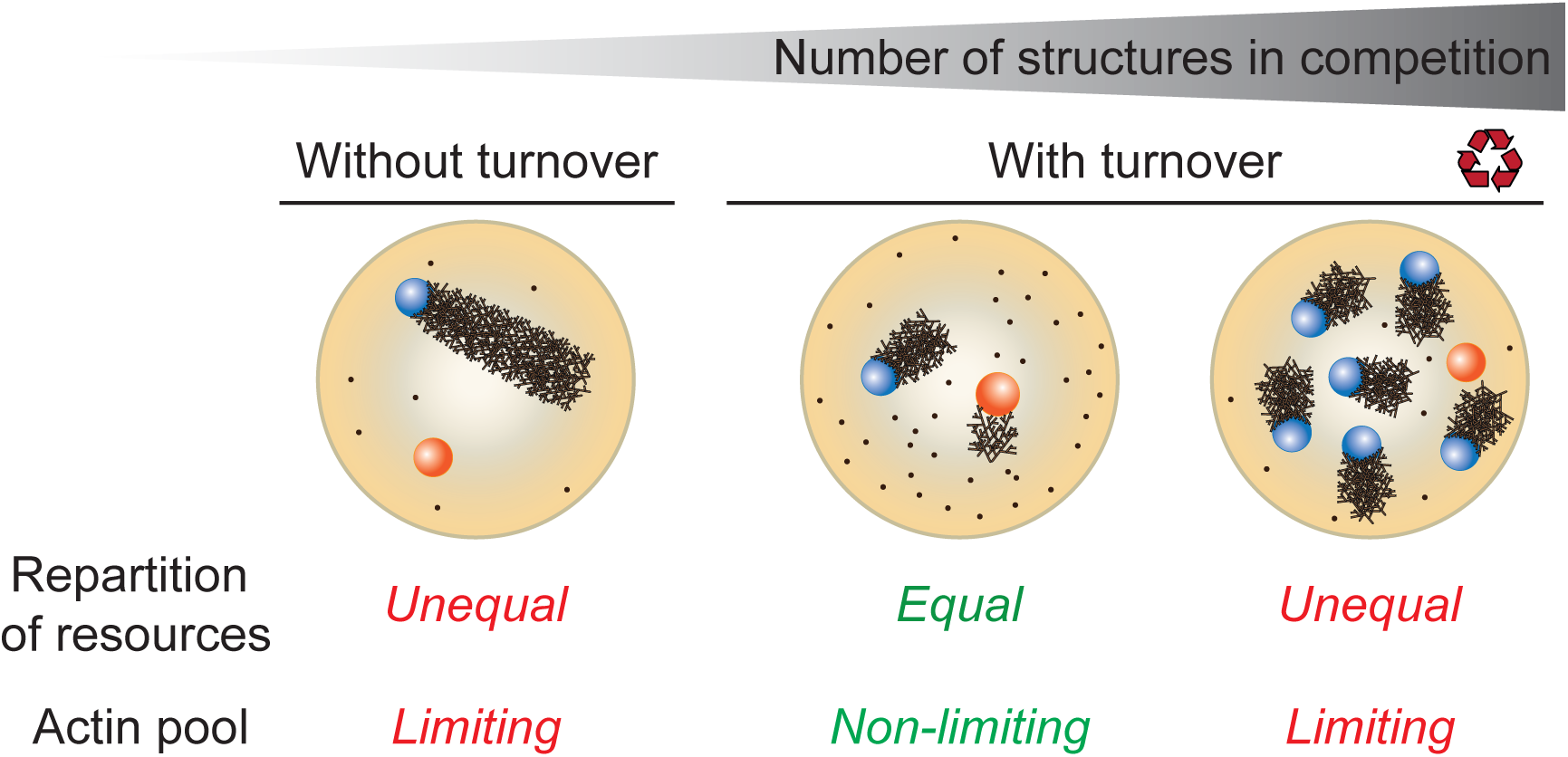

## Introduction

Competition for resources exists at all levels in nature. In ecological communities, the different species sharing the same habitat compete for food resources. Competition for limited resources has been proposed to be a driver of evolution but how to ensure a stable coexistence of various species sharing the same resources is still an open question in ecology (Lankau, 2011; Chesson, 2000). In multicellular organisms, the organs compete for resources during development. Indeed, impeding the growth of one organ will often make the other organs bigger (Gawne et al, 2020; Smiley & Levin, 2022). At a more microscopic level, inter-cellular competition has been proposed to be a surveillance mechanism for the detection and elimination of suboptimal cells during development (Díaz & Moreno, 2005; Merino et al, 2016; Clavería & Torres, 2016). In the tumoral context, cell competition was proposed to have two opposite effects: elimination of pretumoral cells or at later stages, promotion of tumor expansion (Taylor et al, 2017; Moreno, 2008). In those models, the cells compete for resources (trophic factors in the ligand capture model) or for limited space (mechanical constraints). Finally, there is also competition within the cell, due to the finite pool of resources that leads to the budgeting of resources for cell decision-making (Baghdassarian & Lewis, 2024). Indeed, many proteins are shared between several signaling pathways or incorporated into different cellular architectures. This is particularly true for actin, one of the most abundant proteins in the cell, which plays a crucial role in various physiological functions, including maintaining cell architecture, facilitating cell movement, and enabling endocytosis (Blanchoin *et al*, 2014; Lappalainen *et al*, 2022). In recent years, it has become widely accepted that the actin monomer pool is shared among various cellular structures and is limited, thereby influencing notably the size of actin structures (reviewed in (Michelot & Drubin, 2011; Rotty & Bear, 2015; Suarez & Kovar, 2016; Davidson & Wood, 2016; Kadzik et al, 2020)). Indeed, in yeast, Arp2/3 complex inhibition with drugs promotes formin-mediated cable formation, indicating a tension in the actin pool with implications for cell polarity (Sagot et al, 2002; Gao & Bretscher, 2008; Burke et al, 2014; Suarez et al, 2015; Billault-Chaumartin & Martin, 2019). Moreover, multiple studies across various organisms and model systems have shown that actin structures compete within cells, indicating a universal phenomenon where actin homeostasis prevents excessive network outgrowth, such as the Arp2/3 complex regulating formin activity during *C. elegans* embryo cytokinesis (Chan *et al*, 2019). Furthermore, this competition is suggested to impact cell polarization and the initiation of motility (Lomakin *et al*, 2015; Mejillano *et al*, 2004; Dimchev *et al*, 2017, 2021; Cao *et al*, 1992) as well as the size regulation of structures like tunneling nanotubes and microvilli (Henderson *et al*, 2023; Faust *et al*, 2019; Chinowsky *et al*, 2020).

Interestingly, in cells, multiple actin-based structures with various filament density and distinct nucleation rates can coexist despite their competition for actin monomers. Strong nucleators, such as lamellipodial networks, which have a high affinity for monomers that they consume rapidly to elongate numerous filaments in dense arrays, do not prevent the growth of networks from weaker nucleators, such as endocytic buds or cortical networks (Chhabra & Higgs, 2007; Michelot & Drubin, 2011; Campellone *et al*, 2023). It seems that competition does not lead to binary outcomes, such as the survival or death of competitors. How those networks with various strengths can coexist with limited resources is still unknown. A possibility is that the pool of actin monomer is so large that the growth of each network does not restrict the others. However, the various examples above have shown that the growth of high and low consumption networks do depend on each other (Suarez & Kovar, 2016). Another possibility is that the constant disassembly of actin-based structures replenishes the pool and temper the competition. Indeed, most cellular actin architectures turn over within minutes (Goode *et al*, 2023; Théry & Blanchoin, 2024). This turnover allows actin networks to respond to various perturbations or signaling events by adapting the position and size of their various actin-based networks (Plastino & Blanchoin, 2019; Lappalainen et al, 2022; Goode et al, 2023). It might also participate to the co-existence of these networks in a competitive environment for resources. Whether and how actin-based structure turnover modulates their competition for resources and lead to complex outcomes allowing their co-existence, despite limited resources, remains an open question.

Theoretical studies have attempted to uncover mechanisms regulating the size of multiple structures from a finite set of components. Mohapatra *et al*. (Mohapatra et al, 2017) found the limiting pool mechanism insufficient to explain the variation of their sizes. Interestingly, Suarez *et al*. (Suarez et al, 2017) suggested that the various lifetimes of the structures is a key regulator of their competition. More recent works, (Banerjee & Banerjee, 2022b, 2022a) proposed a theory involving negative feedback between growth rate and structure size. However, these theories lack experimental validation, and a comprehensive understanding of how dynamic actin networks share monomer pools remains elusive. Moreover, it is extremely challenging to precisely control the number, position, size and nucleation rates of all actin-based structures in cells. This is why, in this study, we used a reconstituted system to investigate the parameters that enable the coexistence of actin networks in a controlled environment with limited resources. We used the actin-based motility assay to reconstitute branched actin networks (without or with turnover, as described in (Colin *et al*, 2023a)) in microwells in order to limit the enclosed volume and the total amount of monomers. Thereby, we reported the first reconstitution of the competition between dynamic architectures, in the presence of a limited pool of components. We show that the turnover of actin architectures ensures the redistribution of resources among them and thus allows the coexistence of network of different strengths. Finally, we revealed the limits of this balancing system in the case of a large number of competing networks leading to tougher competition.

## Results

### Without turnover, the size of the actin structures is determined by the pool of available monomers

To study competition for actin monomers between multiple networks, we used the well-described bead motility assay (Loisel *et al*, 1999; Cameron *et al*, 1999; Bernheim-Groswasser *et al*, 2002; Cameron *et al*, 2004; Akin & Mullins, 2008; Reymann *et al*, 2011; Kawska *et al*, 2012; Colin *et al*, 2023a). Briefly, we coated 4.5 µm diameter polystyrene beads with a Nucleation-Promoting Factor (NPF, Snap-Streptavidin-WA-His, see Methods) triggering actin assembly via the Arp2/3 complex. We introduced these beads in cell-sized microwells (with a diameter of 100 µm, previously described in (Yamamoto *et al*, 2022; Colin *et al*, 2023a)). The use of these microwells ensures control and limitation of the quantity of proteins present for the growth of actin structures (Kandiyoth & Michelot, 2023). We first grew actin comet tails in presence of actin, profilin, capping protein and Arp2/3 complex (see Methods for concentrations) (**Figure 1A**). In these conditions, the turnover rate is extremely slow (less than 0,02 h^-1^, (Colin *et al*, 2023a)) so we consider this condition as “without turnover”. We imaged the microwells containing the beads with fluorescence microscopy and we analyzed the actin comet size as a function of number of beads in the microwells (**Figure 1B**, **Movie S1**). We calculated structure size from the measurement of comet area (Methods). By measuring comet area per bead as a function of time (**Figure 1C**), we estimated that the growth of comets is completed in about 150 minutes. We therefore decided to measure the area of each individual comet at the growth plateau, between 200 and 260 minutes. When the number of beads in the microwell increases, the size of each individual comet tail decreases, inversely proportional to the number of beads (**Figure 1D**). The estimate of actin polymerized within comet tails (Methods) showed that the quantity of actin polymerized is approximately constant regardless of the number of beads in the microwell (**Figure 1E**). This can also be assessed by quantifying the amount of actin (monomers and oligomers, (Colin *et al*, 2023a)) present in the microwell bulk at steady state, which remains constant regardless of the number of the beads in the microwell (**Figure S1A**). Interestingly, mean density of comet tails slightly decreases when number of beads in the microwell increases (**Figure S1B**), demonstrating that there is probably a competition for nucleation resources when multiple structures grow simultaneously. From these data, we conclude that without turnover, the size of actin structures depends on the number of competing structures. This size variation and the dependency with one over the number of the beads can be attributed to a well-described mechanism involving a limited pool of resources (Goehring & Hyman, 2012; Marshall, 2020), where the actin pool is constrained and distributed equally among the various architectures (**Figure 1F**).

**Figure 1.**
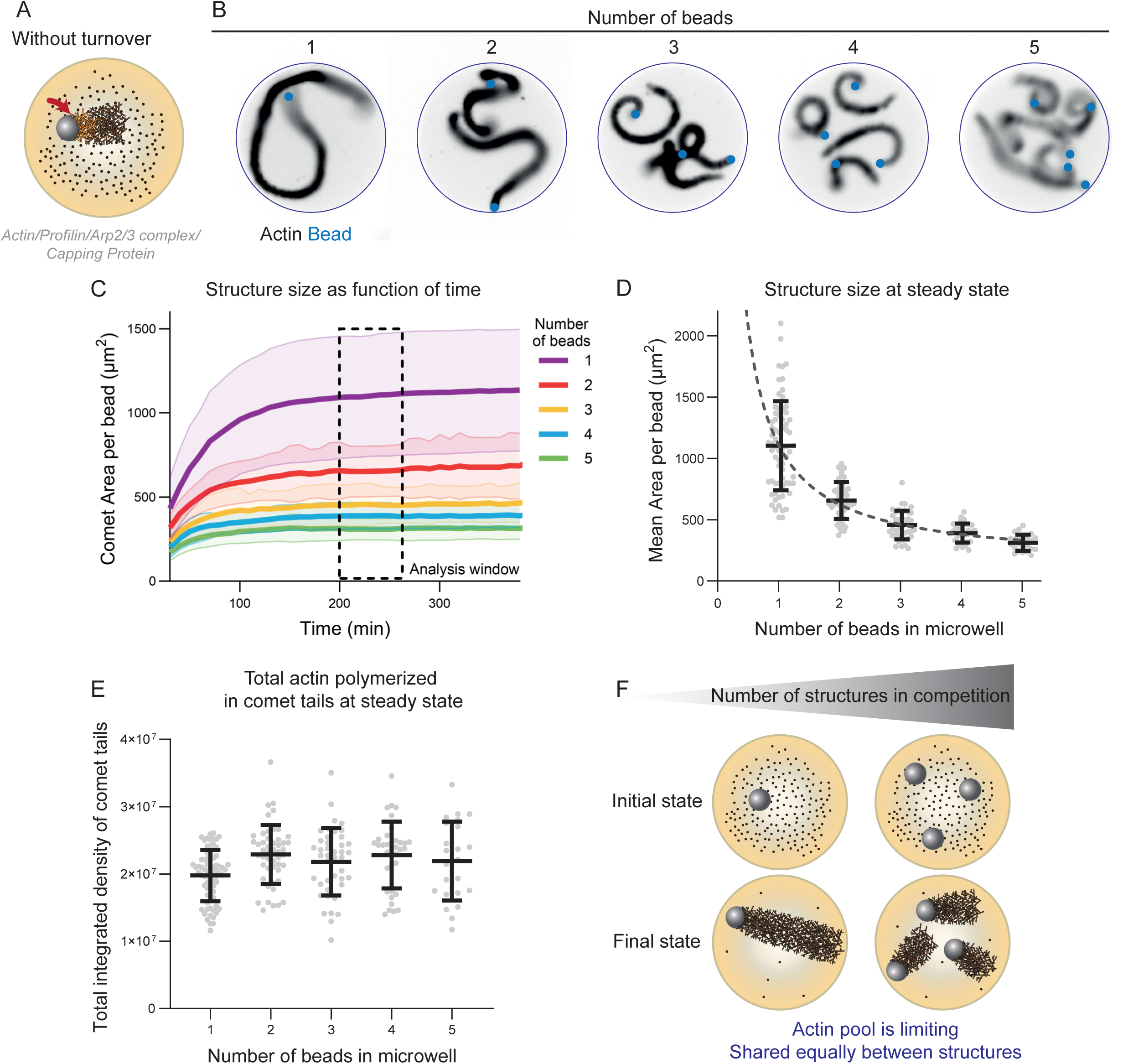
Without turnover, the size of the actin structures is determined by the pool of available monomers. A. Scheme of the experimental set-up (arrow shows the site of the actin network assembly). B. Snapshots of actin comets (black) grown from polystyrene beads (blue) coated with a nucleation-promoting factor (NPF) of the Arp2/3 complex in microwells. Snapshots were taken after 6 hours. C. Mean comet area per bead as a function of time for various number of beads in the microwell. Mean and standard deviation are represented. D. Mean comet area (structure size) per bead at steady state (between 3 and 4 hours) as a function of the number of beads in the microwell. Grey circles represent individual microwells on top of which mean and standard deviation of the whole population are plotted. Dashed line represents a fit (Area = A + B/number of beads, fit results: A = 136.2; B = 976.3, R² = 0.996). E. Total actin quantity in comet tails in steady state as a function of the number of beads in the microwell. Grey circles represent individual microwells on top of which mean and standard deviation of the whole population are plotted. F. Interpretation scheme: without turnover, the network size is determined by the available pool of proteins. <u>Replicates</u>: N = 4 independent experiments. 1 bead: n = 70 microwells, 2 beads = n = 49 microwells, 3 beads: n = 42 microwells, 4 beads: n = 36 microwells, 5 beads: n = 23 microwells. <u>Biochemical conditions</u>: 4.5 µm beads coated with 400 nM of SNAP-Strep-WA-His. Reaction mix: [Actin] = 6 µM. [Profilin] = 12 µM. [Arp2/3 complex] = 90 nM. [Capping Protein] = 40 nM.

### With turnover, the actin monomer pool is partially limited depending on turnover rate and is equally shared between competing structures

Next, we grew comet tails in microwells with slow or rapid actin turnover, incorporating disassembly (ADF/cofilin) and recycling (Cyclase Associated Protein, CAP) proteins at different concentrations (**Figure 2A**), to observe their effect on the size of competitive actin networks. With rapid turnover, the beads move at an average speed of 3 µm.min^-1^ and the comets have an average length of 10 µm, giving a turnover rate of about 0,3 min^-1^. In other words, the comet is renewed every 3.3 minutes, which is similar to the turnover rate observed for the lamellipodia in cells (Wang, 1985; Theriot & Mitchison, 1991; Watanabe & Mitchison, 2002). We considered microwells containing varying number of beads (**Figure 2B**, **Movie S2**). By monitoring the size of actin comets as a function of time (**Figure 2C**), we found that steady state is established after 50 minutes and is stable for over 300 minutes (as previously described in (Colin *et al*, 2023a)). We quantified the comet area at steady state (between 120 and 180 minutes, **Figure 2D**). Remarkably, under conditions of rapid turnover, the size of the comet tails becomes nearly independent of the number of beads in the microwells. Consequently, the total amount of actin polymerized in the comet tails increases with the number of beads in the microwell (**Figure 2E**), whereas it remains constant in conditions without turnover (**Figure 1E**). This demonstrates that rapid turnover dictates the size of the actin comet tails and constrains competition among different actin networks (**Figure 2F**). Interestingly, in slow turnover conditions, we observed that the size of individual comet tails depends on the number of beads in the microwell (**Figure 2D**, blue curve), as observed in conditions without turnover (**Figure 1D**). The total amount of actin polymerized in comet tails still increase with number of beads in the microwell in those conditions (**Figure 2E**), demonstrating that the actin pool is only partially limited. Based on this result, we investigated how limited or rapid turnover could influence the coexistence of branched networks differing in density.

**Figure 2.**
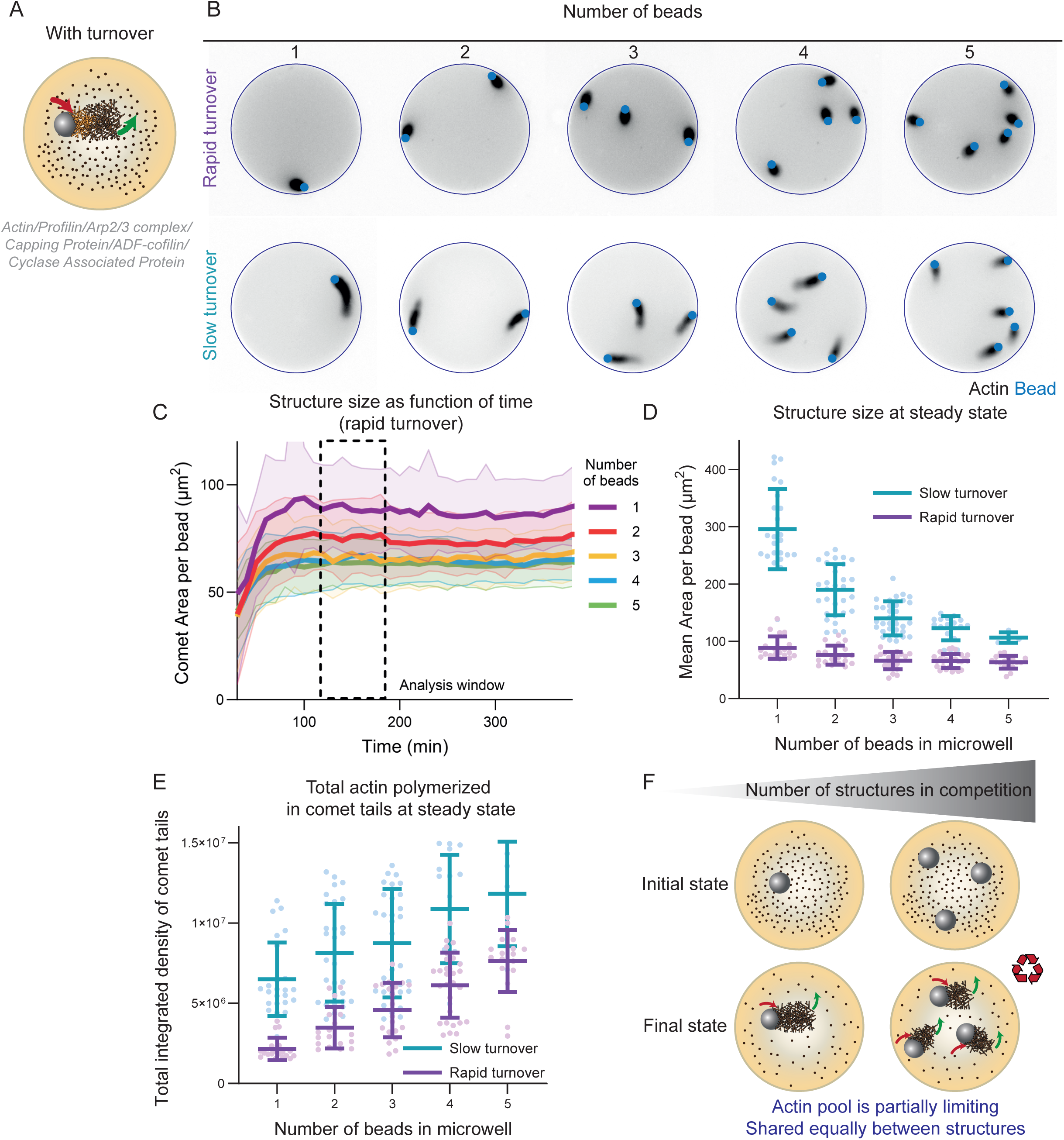
Depending on turnover rate, actin pool is no longer limiting. A. Scheme of the experimental set-up. Red and green arrows indicate assembly/disassembly, respectively. B. Snapshots of actin comets (black) grown from polystyrene beads (blue) coated with a nucleation-promoting factor (NPF) of the Arp2/3 complex in microwells with a high or low disassembly rate (see below for biochemical conditions). Snapshots were taken after 90 minutes. C. Mean comet area per bead as a function of time for various number of beads in the microwell (high disassembly conditions). Mean and standard deviation are represented. D. Mean comet area per bead at steady state (between 2 and 3 hours) as a function of the number of beads in the microwell for low (blue) and high (pink) disassembly rates. Circles represent individual microwells on top of which mean and standard deviation of the whole population are plotted. E. Total actin quantity in comet tails at steady state as a function of the number of beads in the microwell for low (blue) and high (pink) disassembly rates. Circles represent individual microwells on top of which mean and standard deviation of the whole population are plotted. F. Interpretation scheme: depending on turnover rate, actin pool is no longer limiting and is shared equally between the structures. Replicates: Low disassembly rate: N = 2 independent experiments. 1 bead: n = 19 microwells, 2 beads: n = 19 microwells, 3 beads: n = 20 microwells, 4 beads: n = 28 microwells, 5 beads: n = 17 microwells. High disassembly rate: N = 2 independent experiments. 1 bead: n = 24 microwells, 2 beads: n = 31 microwells, 3 beads: n = 34 microwells, 4 beads: n = 17 microwells, 5 beads: n = 4 microwells. Biochemical conditions: 4.5 um beads coated with 400 nM of SNAP-Strep-WA-His. Reaction mix: [Actin] = 6 uM. [Profilin] = 12 uM. [Arp2/3 complex] = 90 nM. [Capping Protein] = 40 nM. High disassembly rate: [ADF/cofilin] = 800 nM. [CAP] = 400 nM. Low disassembly rate: [ADF/cofilin] = 400 nM. [CAP] = 200 nM.

### Without turnover, weak actin networks cannot coexist with strong networks if they are in competition

We prepared two types of beads with different concentrations of NPF (200 nM and 400 nM, referred to as low and high NPF densities, colored orange and blue respectively in **Figures 3, 4 and 5**). The low density beads have a surface density of 3.8 NPF per 100 nm², while the high density beads have a surface density of 7.6 NPF per 100 nm² (similar to the range reported in (Wiesner *et al*, 2003)). When grown in bulk (unlimited resources), the low and high-NPF density beads generated networks with sparse and dense densities respectively (**Figure S2A**). The growth of comet tails with low-NPF density was initiated more slowly (**Figure S2B**, as also reported in Wiesner et al., 2003) and they incorporated actin at a slower rate, as it can be seen from a linear fit on the quantity of actin in the comet tail as a function of time (**Figure S2B**). Overall, given these properties of density and actin consumption rate, we define those networks as weak and strong respectively. We then placed the low and high NPF density beads in microwells. Without competition, both weak and strong actin tails were able to grow (**Figure 3A** Left, **Movie S3**) and reached a maximum size that was similar (**Figure 3B** Left). We then looked at the cases where the two networks (weak and strong) were in competition in the same microwell (**Figure 3A** Right, **Movie S3**). Without turnover, the strong network maintained a size similar to that observed without competition. However, in all cases we tested, the weak network was unable to grow (**Figure 3A, 3B** Right, **Movie S3**). This results in a size decrease of 100% for the weak network, whereas the size of the strong network decreases by only 3% (**Figure 3E** Left). Interestingly, regardless of the number of low NPF density beads in the microwell, their growth cannot initiate, whereas the high-NPF density beads always initiate growth (**Figure S2E, F**). To understand why the weak network cannot initiate growth in the presence of competition, we first examined the kinetics of comet growth in the absence of competition. From the plots depicting the comet area over time (**Figure S2C**), it is evident that there is an approximately 40-minute delay for the weak network (orange curve) to initiate its growth, compared to the strong network. If we analyze the actin available in the microwell bulk (Methods), we observe that by the time the weak network begins to grow (at 40 minutes, **Figure S2D**), the strong network has already consumed 35% of the available actin. This indicates that the weak network encounters only a concentration of ∼3.9 µM of actin, which is insufficient for it to initiate its growth efficiently as observed from the growth of comet tails in flow chamber (**Figure S2G, H**).

**Figure 3.**
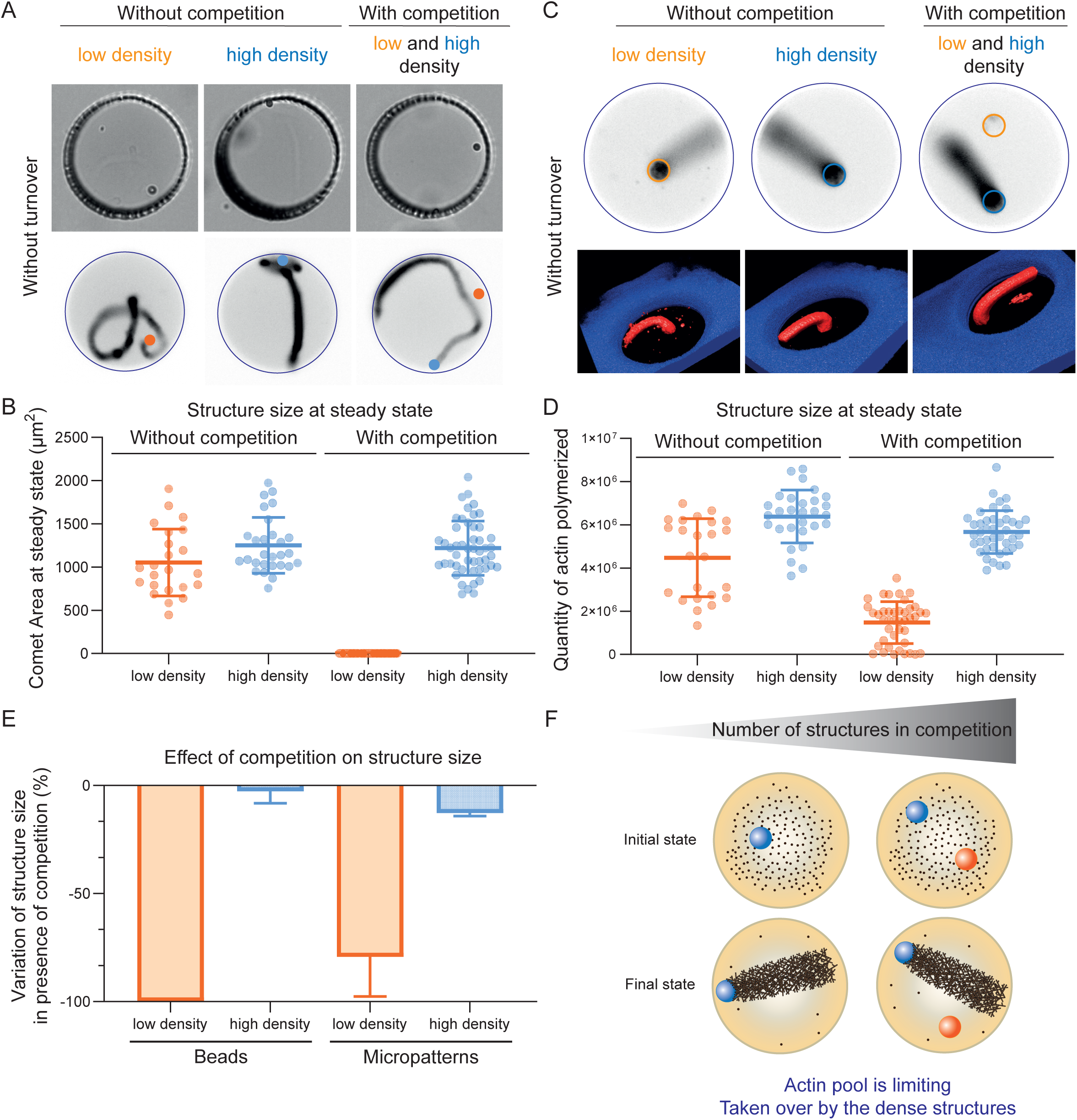
When structures with various densities are in competition without turnover, the actin monomer pool is taken over by dense structures. A. Snapshots (top row: bright field, bottom row: fluorescence with actin in black) of actin comet tails grown from polystyrene beads coated with a nucleation-promoting factor (NPF, blue bead: high-NPF density - 400 nM WA, orange bead: low-NPF density - 200 nM WA) of the Arp2/3 complex in microwells in conditions without turnover. Snapshots were taken after 300 minutes. B. Comet area at steady state for comets grown from low-NPF density (orange) and high-NPF density (blue) beads, in absence or presence of competition. Colored circles represent individual microwells on top of which mean and standard deviation of the whole population are plotted. C. Snapshots (top row) and 3D-reconstructions (bottom row) of actin branched networks grown from lipid micropatterns with various densities of NPF (orange: low and blue: high). D. Quantity of actin polymerized at steady state for structures grown on low-density and high-density patterns, in absence or presence of competition. Colored circles represent individual microwells on top of which mean and standard deviation of the whole population are plotted. E. Variation of structure size in presence of competition expressed as a percentage. The mean variation of structure size is computed per independent dataset and standard deviation represents the variation between independent datasets. F. Interpretation scheme: without turnover, when structures with various densities are in competition, the actin monomer pool is limiting and is taken over by the dense structures. <u>Replicates</u>: Beads: N = 3 independent experiments. Without competition, low-NPF density: n = 23 microwells. Without competition, high-NPF density: n = 28 microwells. With competition: n = 49 microwells. Micropatterns: N = 2 independent experiments. Without competition, low-NPF density: n = 24 microwells. Without competition, high-NPF density: n = 29 microwells. With competition: n = 42 microwells. <u>Biochemical conditions</u>: 4.5 µm beads coated with 400 nM or 200 nM of SNAP-Strep-WA-His. Reaction mix for Assembly conditions: [Actin] = 6 µM. [Profilin] = 12 µM. [Arp2/3 complex] = 90 nM. [Capping Protein] = 40 nM. Micropatterns: Lipids with 0.5% of biotin; coating with 50 nM of SNAP-Strep-WA-His. Same reaction mix as for the beads.

**Figure 4.**
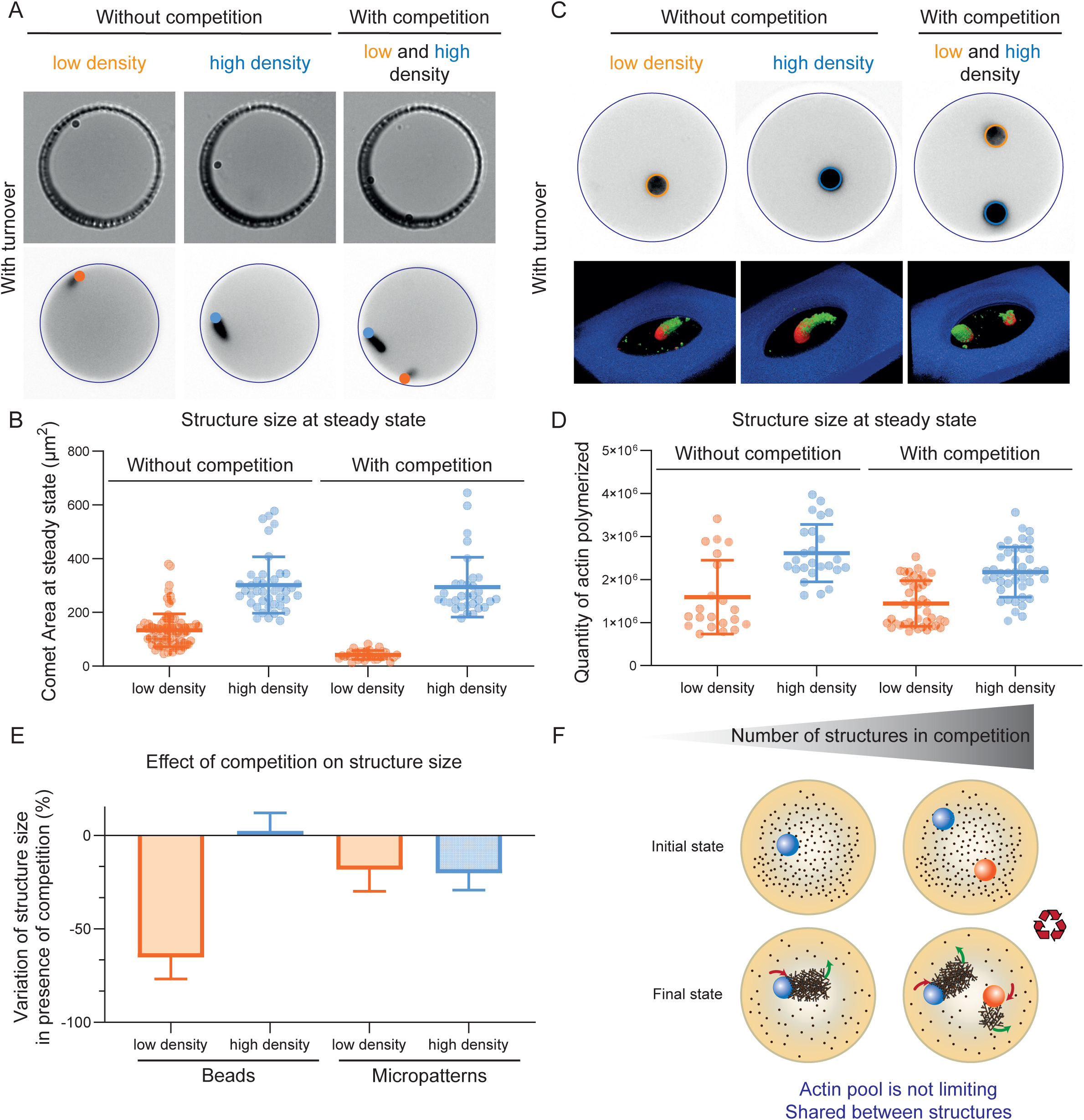
Conditions with turnover allow the coexistence of networks with various densities. A. Snapshots (top row: bright field, bottom row: fluorescence with actin in black) of actin comet tails grown from polystyrene beads coated with a nucleation-promoting factor (NPF, blue bead: high-NPF density - 400 nM WA, orange bead: low-NPF density - 200 nM WA) of the Arp2/3 complex in microwells in conditions with turnover. Snapshots were taken after 240 minutes. B. Comet area at steady state for comets grown from low-density (orange) and high-density (blue) beads, in absence or presence of competition. Colored circles represent individual microwells on top of which mean and standard deviation of the whole population are plotted. C. Snapshots (top row) and 3D-reconstructions (bottom row) of actin branched networks grown from lipid micropatterns with various densities of NPF (orange: weak and blue: strong) in conditions with turnover. D. Quantity of actin polymerized at steady state for structures grown on low-density and high-density patterns, in absence or presence of competition in conditions with turnover. Colored circles represent individual microwells on top of which mean and standard deviation of the whole population are plotted. E. Variation of structure size in presence of competition expressed as a percentage. The mean variation of structure size is computed per independent dataset and standard deviation represents the variation between independent datasets. F. Interpretation scheme: protein turnover is necessary to limit the competition and to allow the coexistence if structures with various densities. <u>Replicates</u>: Beads: N = 3 independent experiments. Without competition, low-NPF density: n = 94 microwells. Without competition, high-NPF density: n = 41 microwells. With competition: n = 32 microwells. Micropatterns: N = 2 independent experiments. Without competition, low-NPF density: n = 21 microwells. Without competition, high-NPF density: n = 25 microwells. With competition: n = 42 microwells. <u>Biochemical conditions</u>: 4.5 µm beads coated with 400 nM or 200 nM of SNAP-Strep-WA-His. [Actin] = 6 µM. [Profilin] = 12 µM. [Arp2/3 complex] = 90 nM. [Capping Protein] = 40 nM. [ADF/cofilin] = 800 nM. [CAP] = 400 nM. Micropatterns: Lipids with 0.5% of biotin; coating with 50 nM of SNAP-Strep-WA-His. [Actin] = 6 µM. [Profilin] = 12 µM. [Arp2/3 complex] = 90 nM. [Capping Protein] = 40 nM. [ADF/cofilin] = 400 nM. [CAP] = 200 nM.

**Figure 5.**
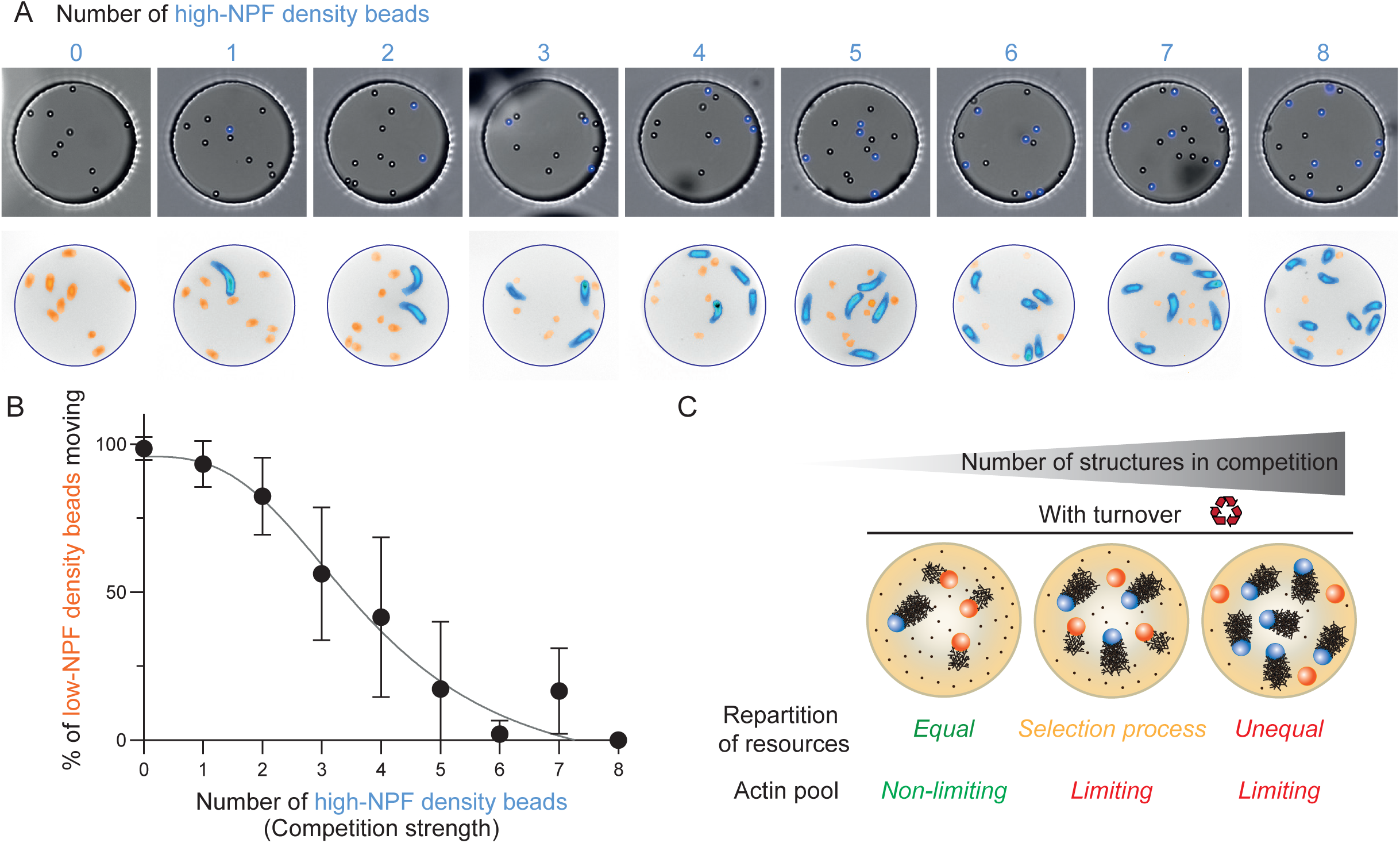
When too many structures are in competition, the actin monomer pool is limiting again. A. Snapshots of microwells with various numbers of high-NPF density beads (blue beads). B. Quantification of the percentage of low-NPF density beads moving (orange beads), as a function of the number of high-NPF density beads (blue beads) in the microwell. C. Interpretation scheme. With turnover, the repartition of resources is equal and the monomer pool is not limiting until a certain number of structures is reached. <u>Biochemical conditions</u>: 4.5 µm beads coated with 400 nM or 200 nM of SNAP-Strep-WA-His. [Actin] = 6 µM. [Profilin] = 12 µM. [Arp2/3 complex] = 90 nM. [Capping Protein] = 40 nM. [ADF/cofilin] = 800 nM. [CAP] = 400 nM. <u>Replicates</u>: N = 2 independent experiments. 0 high-NPF density beads: n = 8 microwells. 1 high-NPF density bead: n = 16 microwells. 2 high-NPF density beads: n = 29 microwells. 3 high-NPF density beads: 23 microwells. 4 high-NPF density beads: 20 microwells. 5 high-NPF density beads: n = 8 microwells. 6 high-NPF density beads: n = 11 microwells. 7 high-NPF density beads: n = 3 microwells. 8 high-NPF density beads: n = 1 microwell.

Since the delay for actin comet growth depends of the initial symmetry-breaking step around the beads, we decided to test if this observation holds in a different geometry where there is no need for symmetry breaking in actin network growth. To do this, we used lipid micropatterns that closely mimic actin polymerization near a membrane, similar to conditions found in cells. We reported the lipid patterning method recently (Colin *et al*, 2023b) and we adapted it here in microwells (see Methods, **Figure S3A**). We passivated the microwells with silanePEG and then used the Primo device (Strale *et al*, 2016; Luo *et al*, 2020) to pattern the selected regions. Following this, we introduced biotin-functionalized lipids and streptavidin-functionalized NPF. Interestingly, using the Primo device, we can adjust the grayscale intensity of the input pattern to vary the degree of PEG layer removal. This results in different quantities of lipids (and consequently NPF) on the micropatterns (**Figure S3B**). The various quantity of NPFs further leads to various actin network density (**Figure S3C**). Therefore, we were able to generate weak and strong actin networks that grew in 3D in the microwell (**Figure 3C**, **Movie S4**). We observed that without turnover, similar to what was described with the beads in competitive scenarios, the weak network could barely grow, whereas it could grow when there was no competition. We confirmed this observation with the 3D reconstruction of the networks and with the quantification of actin polymerized on the different patterns (**Figure 3C, 3D**). In this competitive condition, the size of the weak network decreases by 79%, whereas the strong network size decreases only by 13% (**Figure 3E**). Thus, with another geometry (micropattern), we confirm the results obtained with the beads. We conclude that when there is no turnover, the actin monomer pool acts as a limiting factor for competitive network to coexist. When architectures of different densities compete with each other, the denser networks dominate the available resources in a “winner-takes-all” process, thereby preventing the weak network from growing (**Figure 3F**). We next sought to evaluate whether under conditions with rapid turnover, the system operates differently according to the possible coexistence of network with different strengths.

### Rapid turnover ensures a balanced distribution of resources, thereby enabling the existence of network of different strengths

We then tested conditions with rapid turnover (addition of ADF/cofilin and CAP to the experiment) for beads and micropatterns geometries. For the bead geometry without competition, both weak and strong networks are coexisting, with strong comets a bit longer than weak ones (**Figure 4A, 4B**, **Movie S5**). When the two networks were in competition in the same microwell, we observed that they were both growing and coexisting over a long period of time. The size of the strong network was unaffected by the competition (**Figure 4E**) and the size of the weak network was moderately affected by the competition (decrease of 65%, **Figure 4E**). This shows that rapid turnover enables the existence of weak networks that could not grow without it. Indeed, in presence of turnover, there is a sizeable pool of monomers which is shared because of rapid monomer diffusion. Therefore, some monomers always reach the weak bead surface and can assemble into the tail network there.

We next examined the same configuration with rapid turnover of actin networks grown from micropatterns. Similar to our observations with the beads, we found that in the presence of turnover, both weak and strong networks could grow and coexist (**Figure 4C, 4D**, **Movie S6**). Therefore, we conclude that rapid turnover allows the protein pool to be distributed, accommodating varying strengths of these networks as they grow (**Figure 4F**). However, under competitive condition, both weak and strong networks showed a 20% reduction in size suggesting limitations in this balancing mechanism (**Figure 4E**). We therefore sought to challenge the limitations of the rapid turnover system concerning the coexistence of structures with different strengths.

### When numerous structures compete, the actin monomer pool again becomes limiting, causing the disappearance of the sparse network population

To do this, we examined scenarios where many beads were present in the microwells. We counted the number of high-NPF density beads (in blue on the figure) in the microwell and defined this number as representative of the strength of competition (**Figure 5A**, **Movie S7**). We then examined the percentage of low-NPF density beads (orange on the figure) that initiate growth, as a function of competition strength (**Figure 5B**). We observed that when the level of competition is low (less than two high-density beads), most low-density beads could grow a weak network. Then, in the medium range of competition, we observed a selection process, meaning that only a fraction of the weak network is able to grow and move. When the level of competition is high (more than six high-density beads), weak networks cannot grow (**Figure 5C**). Interestingly, when high-NPF density beads are alone in the microwell (conditions of Figure 2), they reach a plateau for the total actin polymerized in comet tails at steady state after six beads in the microwell (**Figure S4**). We therefore conclude that beyond a certain number of structures, rapid turnover alone cannot ensure the coexistence of all networks, actin pool becomes limiting again and the strongest networks are the ones that ultimately “win it all”.

## Discussion

Our findings highlight the necessity of considering protein turnover to understand the mechanisms that allow the coexistence of various structures competing for a limited pool of components.

Under conditions without turnover, the outcome is dictated by the mere consumption rates of resources and the system is rapidly trapped in its final configuration, preventing further evolution. Weak structures cannot develop, and only strong structures dominate the final architecture. The result is not only homogeneous, but also static. Indeed, while this leads to high stability, it lacks adaptive properties. In cells, some structures like stereocilia or sarcomeres have an extremely low turnover (from hours to days (Manor & Kachar, 2008; Littlefield & Fowler, 2008)). However, both types of structures continue to incorporate actin at their tips, which is necessary for their maintenance through the organism’s life (Belyantseva *et al*, 2009; Perrin *et al*, 2010; Zhang *et al*, 2012; Colpan *et al*, 2021)). An intriguing question is what prevents protein degradation or possible damage in these long-lived structures, as observed in reconstituted systems (Colin *et al*, 2023a). The potential existence of a repair mechanism that maintains cytoskeletal integrity after damage is an exciting possibility (Winkelman *et al*, 2020; Phua *et al*, 2024).

Under turnover conditions, structures begin by consuming resources as they establish a steady-state. Then resources are recycled, and at steady state, the net consumption of resources is effectively zero and the size remains constant. Interestingly, the slow turnover condition represents an intermediate scenario where the slower recycling of resources limits their availability, thereby affecting the size of the structures. Our results also show that competition for resources intensifies when structures with varying strengths coexist. Without turnover, structures deplete resources quickly, leaving little for weaker structures to grow. However, turnover mitigates this issue up to a certain extent. In cells, branched networks involved in key functions like motility and endocytosis compete for actin monomers (Rotty *et al*, 2013; Chakrabarti *et al*, 2021), and within structures like lamellipodia, density variations affect function and movement direction (Bieling *et al*, 2016; Boujemaa-Paterski *et al*, 2017; Manhart *et al*, 2019; Li *et al*, 2022). This study suggests that rapid turnover allows the redistribution of resources and the maintenance of weak structures generated by poor nucleators. Thus, the rapid renewal of dynamic structures such as lamellipodia could allow the coexistence of regions with large variations of densities (Chhabra & Higgs, 2007; Campellone *et al*, 2023). Generally speaking, at all levels of life, turnover is an essential monitoring system and is indispensable for the reuse of components (Reddien, 2024). However, turnover is costly in energy and there is therefore a trade-off between the turnover, the adaptability of the system when facing a perturbation and the energy cost. Reduction or modulation of turnover rate can then be a strategy to reduce the energy consumption, during hibernation (Lewis et al, 2024) or quiescent phases (Sagot et al, 2006) for example.

By examining the system’s limits, we have demonstrated that even under fast turnover excessive competition can force the system to select which structures can polymerize and persist. This decision-making process, which influences the formation or loss of structures, may additionally involve activating signaling pathways or reallocating resources based on turnover rates (Letort *et al*, 2015; Wales *et al*, 2016; Baghdassarian & Lewis, 2024). For instance, cytochalasin-D treatment typically eliminates fast turnover structures, such as lamellipodia, while low-turnover structures, like stress fibers, remain unaffected (Lambert *et al*, 2023). This phenomenon may be attributed to the competition strength caused by cytochalasin-D, which disrupts the coexistence of structures. This work does not address the competition between different actin architectures, such as branched versus linear networks, nor how cellular compartmentalization and heterogeneity might influence this competition (Kadzik *et al*, 2020). Future studies should explore these aspects, including the impact of local actin monomer synthesis (Vedula *et al*, 2021) on the distribution among competing actin networks.

Our study emphasizes the crucial role of competition strength, shaped by turnover rate, protein pool size, and number of competing structures. These factors interact through feedback mechanisms, with high turnover reducing competition and lower turnover increasing protein consumption, therefore affecting the coexistence of actin architectures. Modulating competition strength, by varying the amount of available monomers, significantly affects actin structure size and shape, impacting cellular functions like polarization and migration (Vitriol *et al*, 2013; Liu *et al*, 2015; Lomakin *et al*, 2015; Lee & Kumar, 2020). Additionally, a larger pool of monomers may reduce the competition strength, as evidenced by the increased number of stress fibers in larger cells (Mertz *et al*, 2012; Reinhart-King *et al*, 2005). Interestingly, contrary to the notion that “bigger is better”, evolution appears to favor finite reservoirs, which leads to higher fitness and consistency (Smiley & Levin, 2022). Indeed, a limited protein pool and a high competition strength enable structures to assess each other’s activity based on resource availability, with protein concentration influencing turnover rates and assembly (Rafelski & Marshall, 2008; Goehring & Hyman, 2012; Gawne *et al*, 2020; Smiley & Levin, 2022).

## Acknowledgments

We thank Benoit Vianay for insightful discussions. This work was supported by the European Research Council (Consolidator Grant 771599 (ICEBERG) to MT and Advanced Grant 741773 (AAA) to LB). This work was also supported by the MuLife imaging facility, which is funded by GRAL, a program from the Chemistry Biology Health Graduate School of University Grenoble Alpes (ANR-17-EURE-0003).

## Author contributions

Design of research: CG, AC, LB, MT, AM / Experiments: AC, CG / Preliminary observations: ABN, LG / Analysis: AC / Funding acquisition: LB, MT / Writing of the paper: AC, LB, MT

## Movies legends

**Movie S1 – Actin comet tails assembly in microwells without turnover**

Movie corresponding to snapshots in Figure 1B. Time-lapse of various number of beads in microwells for comets grown in conditions without turnover. Bead is shown in blue and labelled actin in black. 1 image was taken every 10 minutes during 6 hours. The movie was compressed in JPEG at 5 frames per seconds.

**Movie S2 – Actin comet tails assembly in microwells with rapid turnover**

Movie corresponding to snapshots in Figure 2B. Time-lapse of various number of beads in microwells. Comets grown in conditions with turnover. Bead is shown in blue and labelled actin in black. 1 image was taken every 10 minutes during 6 hours. The movie was compressed in JPEG at 5 frames per seconds.

**Movie S3 – Actin comet tails with various densities in microwells without turnover**

Movie corresponding to snapshots in Figure 3A. Time-lapse of beads with various densities without or with competition in conditions without turnover. High-NPF density bead is shown in blue, low-NPF density bead is shown in orange and labelled actin in black. 1 image was taken every 10 minutes during 6 hours. The movie was compressed in JPEG at 7 frames per seconds.

**Movie S4 – Actin networks grown from micropatterns with various densities in microwells without turnover**

Movie corresponding to snapshots in Figure 3C. Time-lapse of actin networks grown from micropatterns coated with various densities of NPF without or with competition in conditions without turnover. High-NPF density pattern is shown in blue, low-NPF density pattern is shown in orange and labelled actin in black. 1 image was taken every 5 minutes during 140 minutes. The movie was compressed in JPEG at 7 frames per seconds.

**Movie S5 – Actin comet tails with various densities in microwells with turnover**

Movie corresponding to snapshots in Figure 4A. Time-lapse of beads with various densities without or with competition in conditions with turnover. High-NPF density bead is shown in blue, low-NPF density bead is shown in orange and labelled actin in black. 1 image was taken every 10 minutes during 6 hours. The movie was compressed in JPEG at 7 frames per seconds.

**Movie S6 – Actin networks grown from micropatterns with various densities in microwells with turnover**

Movie corresponding to snapshots in Figure 4C. Time-lapse of actin networks grown from micropatterns coated with various densities of NPF without or with competition in conditions with turnover. High-NPF density pattern is shown in blue, low-NPF density pattern is shown in orange and labelled actin in black. 1 image was taken every 5 minutes during 140 minutes. The movie was compressed in JPEG at 7 frames per seconds.

**Movie S7 – High number of beads with various densities in microwells with turnover**

Movie with a selection of snapshots from Figure 5A. Time-lapse of beads with various densities in conditions with turnover. Strong density bead is shown in blue, weak density bead is shown in orange and labelled actin in black. 1 image was taken every 10 minutes during 280 minutes. The movie was compressed in JPEG at 7 frames per seconds.

## Materials and Methods

### Protein Purification

Actin was purified from rabbit skeletal-muscle acetone powder (Spudich & Watt, 1971). Monomeric Ca-ATP-actin was purified by gel-filtration chromatography on Sephacryl S-300 at 4°C in G buffer (2 mM Tris–HCl, pH 8.0, 0.2 mM ATP, 0.1 mM CaCl_2_, 1 mM NaN_3_ and 0.5 mM dithiothreitol (DTT)). Actin was labelled on lysines with Alexa-568 (Isambert *et al*, 1995). All experiments were carried out with 5% labelled actin. The Arp2/3 complex was purified from calf thymus according to (Egile *et al*, 1999) with the following modifications: the calf thymus was first mixed in an extraction buffer (20 mM Tris pH 7.5, 25 mM KCl, 1 mM MgCL2, 0.5 mM EDTA, 5% glycerol, 1 mM DTT, 0.2 mM ATP and proteases). Then, it was placed in a 50% ammonium sulfate solution in order to make the proteins precipitate. The pellet was resuspended in extraction buffer and dialyzed overnight.

Human profilin was expressed in BL21 DE3 pLys *Escherichia coli* cells and purified according to (Almo *et al*, 1994). Mouse capping protein was purified according to (Palmgren *et al*, 2001). Yeast cofilin was purified and fluorescently-labelled according to (Suarez *et al*, 2011). The full-length mouse cyclase associated protein 1 (CAP1) was purified in a similar fashion as described in (Colin *et al*, 2023a). Snap-Streptavidin-WA-His was purified as described in (Colin *et al*, 2023a).

### Polystyrene Beads Coating

Polystyrene beads coating with NPF was done following classical protocols ((Reymann *et al*, 2011)). 20 µL of 4.5 µm polystyrene beads (Fluoresbrite BB Carboxylate Microspheres, 2.5% solids) were centrifuged at 13, 000 g for 2 minutes on a mini spin plus Eppendorf centrifuge (Rotor F45-12-11). The pellet was then resuspended in 200 µL of a 400 nM or 200 nM Snap-Streptavidin-WA-His solution. Beads were incubated for 15 minutes at 15°C at 950 rpm in a thermoshaker. They were then centrifuged for 2 minutes at 3800 g, resuspended in 200 µL of BSA 1% and let on ice for 5 minutes. Beads were finally centrifuged again 2 minutes at 3800 g and resuspended in 50 µL of BSA 0.1%. The bead coating was redone every day.

<u>Estimation of protein density on beads</u>: for a coating with 400 nM of NPF, we estimate that we have about 7,6 NPF molecule per 100 nm^2^. Therefore, for a coating with 200 nM of NPF, this represents about 3,8 NPF molecule per 100 nm^2^. These number are in the same order of magnitude as the conditions used in (Wiesner *et al*, 2003).

### Microwells Preparation

SU8 mold with Pillars was prepared using standard protocols and silanized with Trichloro(1H,1H,2H,2H-perfluoro-octyl)silane for 1 hour and heated for 1 hour at 120°C. From the SU8 mold, a PDMS primary mold was prepared (Dow, SYLGARD 184 silicone elastomer kit) with a 1:10 w/w ratio of curing agent. PDMS was cured at 70°C for at least 2 hours. PDMS primary mold was then silanized with Trichloro(1H,1H,2H,2H-perfluoro-octyl)silane for 1 hour and heated for 2 hours at 100°C. PDMS was then poured on top of the PDMS primary mold to prepare the PDMS stamps.

Coverslips were cleaned with the following protocol: they were first wiped with ethanol (96%) then sonicated 30 minutes in Hellmanex 2% at 60°C. After the sonication, coverslips were rinsed in several volumes of mqH_2_O and kept in water until use. Just before use, coverslips were dried with compressed air.

For the microwells preparation, PDMS Stamps were cut in pieces and placed on the coverslips with the pillars facing the coverslip. A drop (5 µL) of NOA 81 (Norland Products) was then placed on the side of the PDMS stamp and NOA was allowed to go through the PDMS Stamp by capillarity. When the NOA filled all the stamp, it was polymerized with UV light for 12 minutes (UV KUB2/ KLOE; 100% power). After polymerization of the NOA, PDMS stamp was removed and the excess of NOA was cut. Then, an additional UV exposure of 2 minutes was done and the microwells were placed on a hot plate at 110°C for at least 3 hours (or at 60°C overnight) to tightly bind the NOA to the glass.

### Lipids/SUV preparation

For regular microwell passivation, L-α-phosphatidylcholine (EggPC) (Avanti, 840051C, 10 mg/mL) was used. The SUV were prepare as following: 100 µL of Lipids were introduced in a glass tube and the mixture was dried with nitrogen gas. The dried lipids were incubated in a vacuum overnight. After that, the lipids were hydrated in the SUV buffer (10 mM Tris (pH 7.4), 150 mM NaCl, 2 mM CaCl_2_). The mixture was sonicated on ice for 10 minutes. The mixture was then centrifuged for 10 min at 20,238 x g to remove large structures. The supernatants were collected and stored at 4°C.

For the lipids used for the micropatterning, same protocol was used. The lipid mix was composed of L-α-phosphatidylcholine (EggPC) (Avanti, 840051C), DSPE-PEG(2000) Biotin (1,2-distearoyl-sn-glycero-3 phosphoethanolamine-N-[biotinyl(polyethylene glycol)-2000], ammonium salt, Avanti,: 880129C-10mg chloroform) and ATTO 390 labeled DOPE (ATTO-TEC, AD 390-161 dehydrated). Lipids were mixed in glass tubes as follows: 98.5% EggPC (10 mg/mL), 0.5% DSPE-PEG(2000) Biotin (10 mg/mL) and 1% DOPE-ATTO390 (1 mg/mL).

### SilanePEG30k Slides

SilanePEG (30kDa, PSB-2014, Creative PEG works) was prepared at a final concentration of 1 mg/mL in 96% ethanol and 0.1%(v/v) HCl. Slides were cleaned with the following protocol: they were sonicated for 30 minutes at 60°C in Hellmanex 2%. They were then rinsed with several volumes of mqH_2_O and dried with compressed air. Slides were then plasma cleaned for 5 minutes at 80% power and directly immersed in the silanePEG solution. They were kept in the silanePEG solution until use.

### Bead Motility assay in microwells

A typical experiment of bead motility in microwells was performed as follows. The coverslip with microwells was activated with plasma for 2 minutes at 80% power. Just after the plasma, the flow chamber was mounted with the microwells coverslip, a slide passivated with SilanePEG 30k and 180 µm height double-side tape. Lipids were then inserted in the flow chamber and incubate for 10 minutes. Lipids were then rinsed with 1 mL of SUV buffer and 200 µL of 1x HKEM buffer (50 mM KCl, 15 mM HEPES pH=7.5, 5 mM MgCl_2_, 1 mM EGTA). The reaction mix with the different proteins was then prepared and flowed in the flow cell.

A typical reaction mix (60 µL) was prepared with beads coated with WA (activator of the Arp2/3 complex) and 6 µM of actin monomers, 12 µM profilin, 90 nM Arp2/3 complex, 40 nM Capping Protein in HKEM Buffer and was supplemented with 0.7% BSA, 0.2% methylcellulose, 2.7 mM ATP, 5 mM DTT, 0.2 mM DABCO (motility buffer). When needed the polymerization mix was also supplemented with yeast cofilin and/or cyclase associated protein (CAP). The microwells were then closed with mineral oil (Paragon scientific Viscosity Reference Standard RTM13). The whole flow cell was then sealed with VALAP and directly imaged under the microscope.

### Micropatterning in microwells

Micropatterning in microwells is done with the Primo device (Alveole). Prior to micropatterning, the microwells were activated with plasma for 5 minutes at 80% power and directly immersed in a SilanePEG (30k, cf section above) solution overnight. They were rinsed with ethanol and water just before used. A flow chamber was then built with the microwells and a slide passivated with SilanePEG with a double-side tape of 250 µm. Liquid photo initiator (PLPP) was then introduced in the flow chamber. Patterns were designed in Inkscape and loaded into the µmanager’s Leonardo plugin (Alveole). The burning was done at 90 mJ at 100% of a 5.6 mW 365 nm wavelength laser. The flow chamber was then rinsed with 1 mL of water and 400 µL of SUV Buffer to remove the PLPP. 80 µL of SUV solution was then added and incubated for 10 minutes at room temperature for an effective lipid coating. The excess of SUV was washed out by passing 400 µL of SUV Buffer and 200 µL of 1x HKEM. The flow chamber was then passivated with 200 µL of 0,1% BSA diluted in 1x HKEM for 2 minutes and then rinsed with 400 µL of 1x HKEM. The NPF (Snap-Streptavidin-WA-His, 50 nM diluted in 1x HKEM and 0,1% BSA) was then injected in the flow chamber and the coating was done by letting incubate for 10 minutes at room temperature. The excess of NPFs was washed out by passing 1 mL of 1x HKEM. The reaction mix was then introduced (60 µL) and the microwells were subsequently closed with oil.

### Imaging

All the experiments were done with an epifluorescence system (Ti2 Nikon inverted microscope equipped with a Hamamatsu Orca Flash 4.0 LT Plus Camera. The following objectives were used: Plan Fluor 10X DIC and S Plan Fluor ELWD 20X DIC. Time lapse were acquired with the NIS elements software (version 4.60).

Z-stacks of actin networks grown on micropatterns in microwells were performed with a confocal spinning-disc system (EclipseTi-E Nikon inverted microscope equipped with a CSUX1-A1 Yokogawa confocal head, an Evolve EMCCD camera (Photometrics), and a Plan Fluor 40X objective). Z-stacks were acquired with Metamorph software (Universal Imaging). Three-dimensional reconstructions were performed with the UCSF Chimera package. Chimera is developed by the Resource for Biocomputing, Visualization, and Informatics at the University of California (San Francisco) with support from the National Institutes of Health (National Center for Research Resources grant 2P41RR001081, National Institute of General Medical Sciences grant 9P41GM103311).

### Image analysis

Images were analyzed with customed macros written in FiJi (Schindelin *et al*, 2012). Data were then processed with R software and plotted with GraphPad Prism. Mean and standard deviation are represented for all the data. The dot plots show the individual values with the mean and standard deviation superimposed. Fits were done with GraphPad Prism.

Actin comets were detected with the following procedure. First, threshold was adjusted manually and images were binarized. Actin comets were detected with the Analyze particles function. Comet length was obtained with the “skeletonize” and “analyze skeleton” functions. Tracking of comets and fluorescent beads was then done with the TrackMate plugin using the thresholding or LoG detectors respectively ((Ershov *et al*, 2022)). In order to reduce experimental noise, the various quantities measured were averaged over one hour. For conditions without turnover, they were averaged between 200 and 260 minutes (dashed box, Figure 1C). For conditions with turnover, they were averaged between 120 and 180 minutes (dashed box, Figure 2C).

#### Estimation of the ratio of actin consumed from bulk in the comet tail

To estimate the total quantity of actin in the microwell, we estimated the total fluorescence intensity. This value was constant during the time course of an experiment showing that the actin in the microwell is constant during an experiment. The quantity of actin inside a comet tail was estimated after thresholding and binarization of the comet. Those two fluorescence intensities were corrected for the fluorescence background. Then, we estimated the ratio of actin consumed from the bulk in the comet by computing the comet fluorescence over the total fluorescence of the microwell.

---

**Figure S1:**
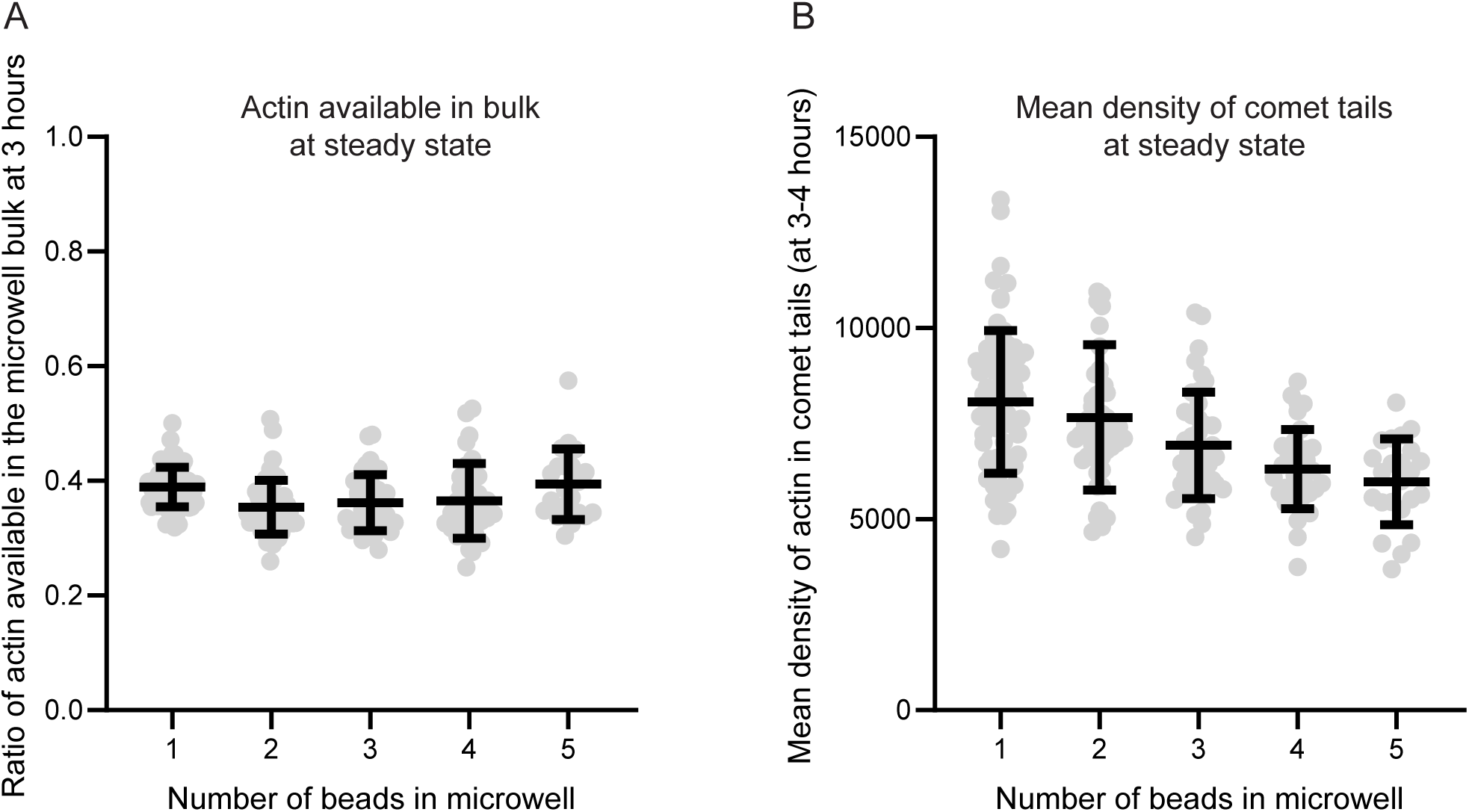
Without turnover actin pool is limiting (related to Figure 1). A. Ratio of actin available in the microwell bulk at steady state. Grey circles represent individual microwells on top of which mean and standard deviation of the whole population are plotted. B. Mean density of actin in comet tails at steady state as a function of the number of beads in the microwell. Grey circles represent individual microwells on top of which mean and standard deviation of the whole population are plotted.

**Figure S2:**
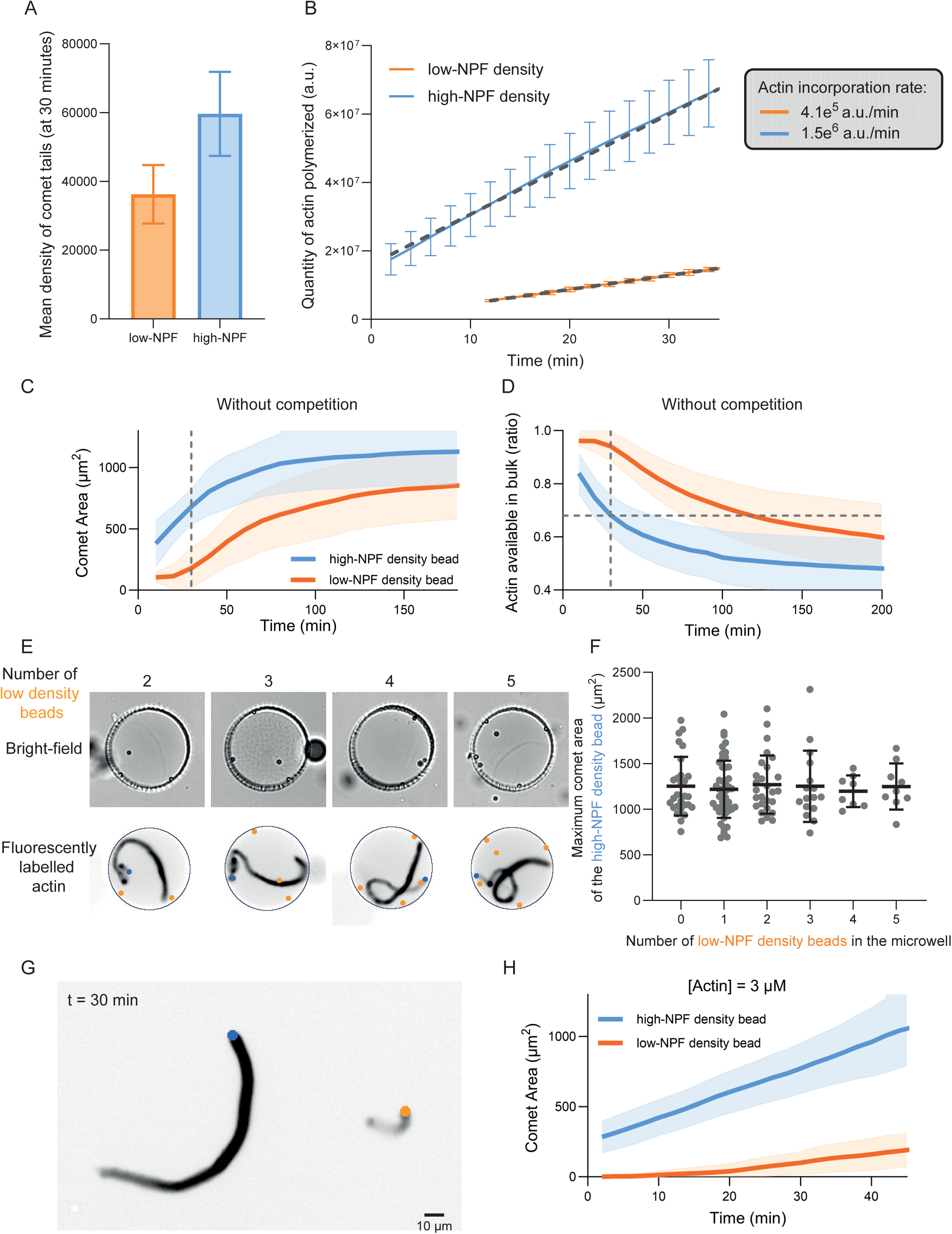
Low-NPF density beads cannot start their growth because resources are taken over by high-NPF density beads (related to Figure 3). A. Mean density of comet tails measured at 30 minutes for comets grown in flow-chamber (unlimited resources). B. Quantity of actin polymerized in comet tails as a function of time for comets grown in flow-chamber (unlimited resources). Dashed line represents a linear fit for which slopes values are shown in the grey box. C. Mean comet area (blue: high-NPF density bead, orange: low-NPF density bead) as a function of time in the absence of competition in the microwell. Mean and standard deviation are represented. D. Mean ratio of actin available in the microwell bulk (blue: high-NPF density bead, orange: low-NPF density bead) as a function of time in the absence of competition in the microwell. Mean and standard deviation are represented. E. Snapshots (top row: bright field, bottom row: fluorescence with actin in black) of actin comet tails grown from polystyrene beads coated with a nucleation-promoting factor (NPF, blue bead: high-density, orange bead: low-density) of the Arp2/3 complex in microwells in conditions without turnover. Snapshots were taken after 300 minutes. F. Quantification of the maximum comet area of the high-NPF density bead as a function of various number of low-NPF density beads in the microwell. G. Snapshot of comet tails grown in flow chamber from high-NPF density bead (blue) or low-NPF density bead (orange) after 30 minutes. H. Quantification of comet area as a function of time for comet tails grown from beads with various NPF densities with 3 µM of actin in flow chamber.

**Figure S3:**
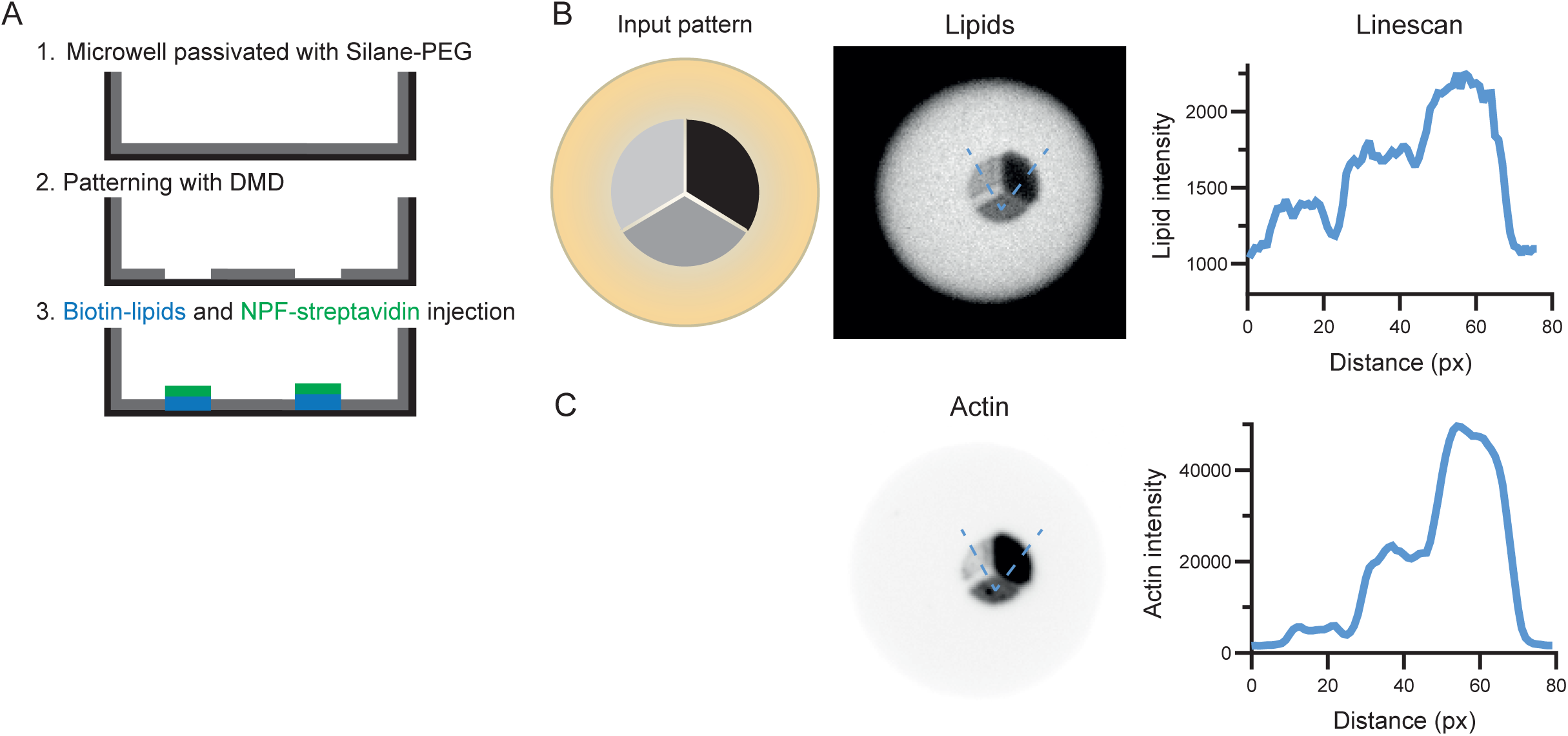
Method for the design of lipid micropatterns with various densities of NPF in microwells (related to Figure 3). A. Scheme of the principle of micropatterning in microwells. B. Proof of principle to generate actin networks with various densities on lipid micropatterns. The input pattern is designed with 3 different grey levels. Depending on the grey levels, there will be more or less lipids on the pattern (right side, image and linescan). C. Resulting actin network: the density of actin network depends on the density of lipids (and therefore NPF) on the micropattern.

**Figure S4:**
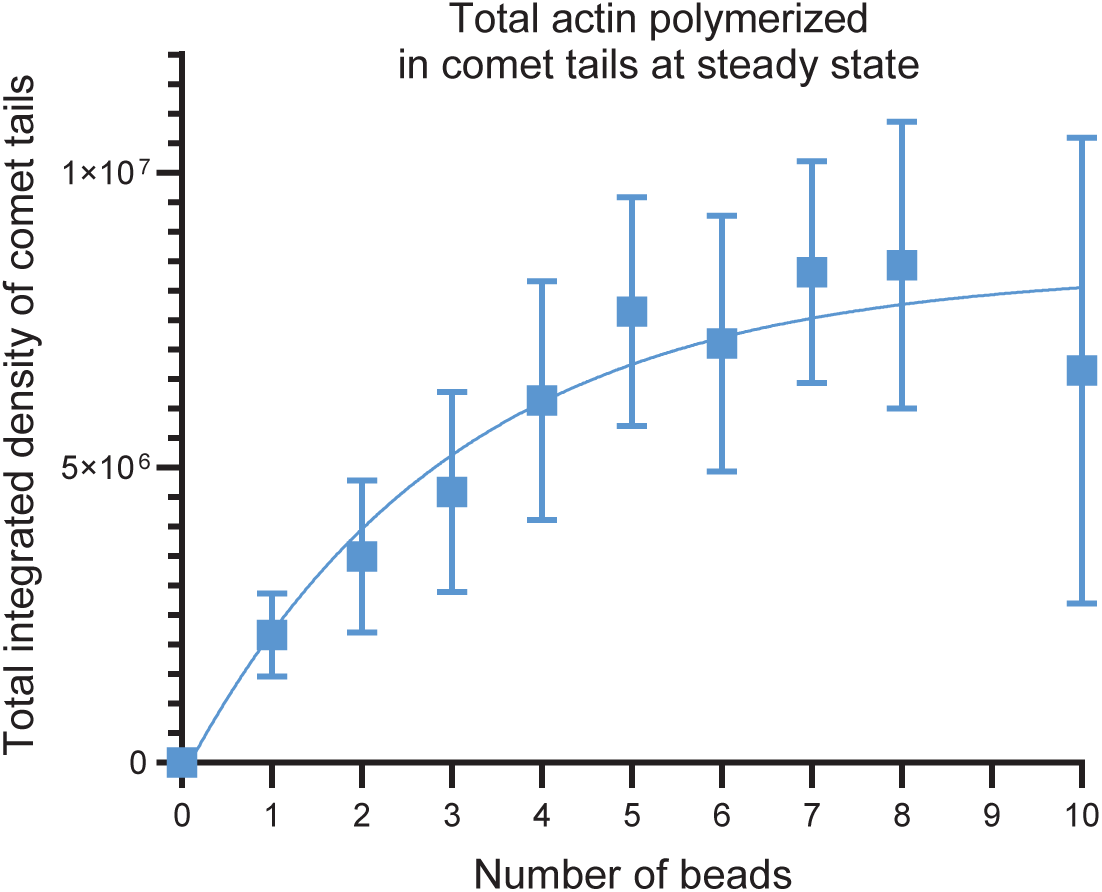
High-NPF density beads reach a plateau for the total actin polymerized in comet tails at steady state after six beads in the microwell (related to Figure 5). Total actin quantity in comet tails at steady state as a function of the number of beads in the microwell with rapid turnover. Mean and standard deviation of the whole population are plotted.

